# Fatiguing Effects of Indirect Vibration Stimulation in Upper Limb Muscles- pre, post and during Isometric Contractions Superimposed on Upper Limb Vibration

**DOI:** 10.1101/429407

**Authors:** Amit N. Pujari, Richard D. Neilson, Marco Cardinale

## Abstract

**Background:** Indirect vibration stimulation i.e. whole body vibration and upper limb vibration (ULV), are gaining popularity as exercise intervention for sports and rehabilitation applications. However, the fatiguing effects of indirect vibration stimulation are not yet fully understood. In addition, current vibration stimulation devices have a series of limitations. For this scope, we investigated the effects of ULV superimposed on fatiguing graded isometric contractions using a newly, purpose developed upper limb stimulation device. Twelve healthy volunteers were exposed to both ULV superimposed to fatiguing isometric contractions, at 80% of the maximum voluntary contractions (V) and just isometric contractions performed on a custom designed arm curl/flexion device- Control (C). The stimulation used consisted of 30Hz frequency of 0.4mm amplitude. Surface electromyographic (EMG) activity of the Biceps Brachii (BB), Triceps Brachii (TB), and Flexor Carpi Radialis (Forearm- FCR) were measured during both V and C conditions. EMG amplitude (EMGrms) and mean frequency (MEF) were computed to quantify muscle activity and fatigue levels respectively.

**Results:** All three muscles BB, TB and FCR displayed significantly higher reduction in MEFs and corresponding significant increase in EMGrms with the V than the C, during fatiguing contractions (P < 0.05). Post vibration treatment, all muscles showed higher levels of MEFs after recovery compared to the control.

**Conclusions:** Our results show that near maximal (80% of MVC) isometric fatiguing contractions superimposed on vibration stimulation lead to a higher rate of fatigue development compared to the isometric contraction alone in the upper limb muscles. Results also show, higher manifestation of mechanical fatigue post treatment with vibration compared to the control. Vibration superimposed on isometric contraction not only seems to alter the neuromuscular function during fatiguing efforts by inducing higher neuromuscular load but also post vibration treatment, potentially through the augmentation of stretch reflex and/or higher central motor command excitability.

## Background

Vibration stimulation has been used as a diagnostic tool in neurological studies since the 70s [1]. However, in recent years, so called ‘indirect vibration’ has been increasingly investigated for its positive effects. Various studies have proposed the use of vibration stimulation for increasing muscle strength, muscle power, balance and bone remodeling [2]–[6]. Consequently, given the potential benefits of vibration stimulation, it has been suggested that this therapeutic modality could represent a viable intervention to improves sport performance and/or enhance rehabilitation from injury [7], [8].

While the potential beneficial effects of vibration stimulation on muscle and bone form and functions are recognized now in various populations [9], [10], a lack of consensus exists on the biological mechanisms responsible for such adaptations. In particular, when assessing the typical neuromuscular responses observed, a muscle tuning hypothesis was suggested [11]. The main idea behind this approach was that the observed increase in neuromuscular activity observed during vibration is a strategy of working muscles to damp the vibratory stimulation [12] and such activity is modulated by sensory receptors [13]. Exposure to such stimulation can therefore enhance neuromuscular function due to the stimulation of sensory motor pathways, leading to an increase in force generating capacity in skeletal muscle.

We have recently developed a portable vibration stimulation device for the upper limbs [14]; this novel and portable device enables the user to perform isometric contraction exercise(s) of various intensities and superimpose a vibratory stimulation. To the best of the authors’ knowledge, only few studies have investigated the effects of graded isometric contractions superimposed on vibration stimulation in the upper limbs [15]. The device developed by Mischi et al. [15]–[17] is capable of superimposing vibration to various levels of muscle contraction by means of a pulley system with an operating frequency of 0–60Hz with a limited pulling force. With this approach, it was shown that the biceps and triceps muscles benefit show an increased EMG activity when vibration is superimposed to force production [17] and that this modality of exercise determines a higher degree of neuromuscular fatigue [15].

However, considering the size of the equipment and the limited operating envelope, it would be beneficial to find a technical solution to provide vibration exercise with smaller equipment and a wider operating envelope. For this reason, and to overcome the limitations of other devices [14] we aimed to develop a smaller vibratory exercise device for the upper limbs and assess its feasibility as a means to increase neuromuscular performance in healthy individuals.

Furthermore, we aimed to assess the effects of vibratory stimulation with this device on higher level of muscle tension to determine the typical neuromuscular responses and identify the possibility to induce fatigue in the target muscles to a larger degree than muscle tension alone.

We hypothesized that compared to the Control (C) condition:

1. The Vibration stimulation (V) would induce an increased neuromuscular activity in the Biceps Brachii (BB), Triceps Brachii (TB), and Flexor Carpi Radialis (Forearm- FCR) muscles as measured by surface EMG
2. V stimulation would induce a higher degree of neuromuscular fatigue as compared to C

## Methods

### Participants

7 female and 6 male (Age 28 years ± 7.24, Height 173 cm± 13.04, Weight 73.16 Kg± 11.19) healthy volunteer participants were recruited through the University of Aberdeen, Biomedical Engineering laboratory. The level of physical training of the participants varied from sedentary to amateur athlete. Written and informed consent was signed for by each volunteer. Experiments were approved by the College Ethics Review Board (CERB) of the College of Life Sciences and Medicine, University of Aberdeen, Aberdeen, Scotland, UK.

Exclusion criteria included history of heart disease and thrombosis, recent musculoskeletal injury, recent fractures in the upper limbs, metallic plates in the bones of the upper limbs, pacemakers, circulatory diseases.

### Experimental setup

Trials were performed with the set-up previously reported in detail [14], [18].

Briefly, the participants sat on the chair with back and arm rest with their arm flexed at a 90° elbow angle while pushing against a vibrating hand grip (Figure 1). Participants tightly gripped the handle bar tightly while exercising. All the participants received detailed instructions on the posture and relevant exercise protocol. It was ensured that there was no contact between participants’ elbow and the chair handle (arm rest).

**Figure 1:**
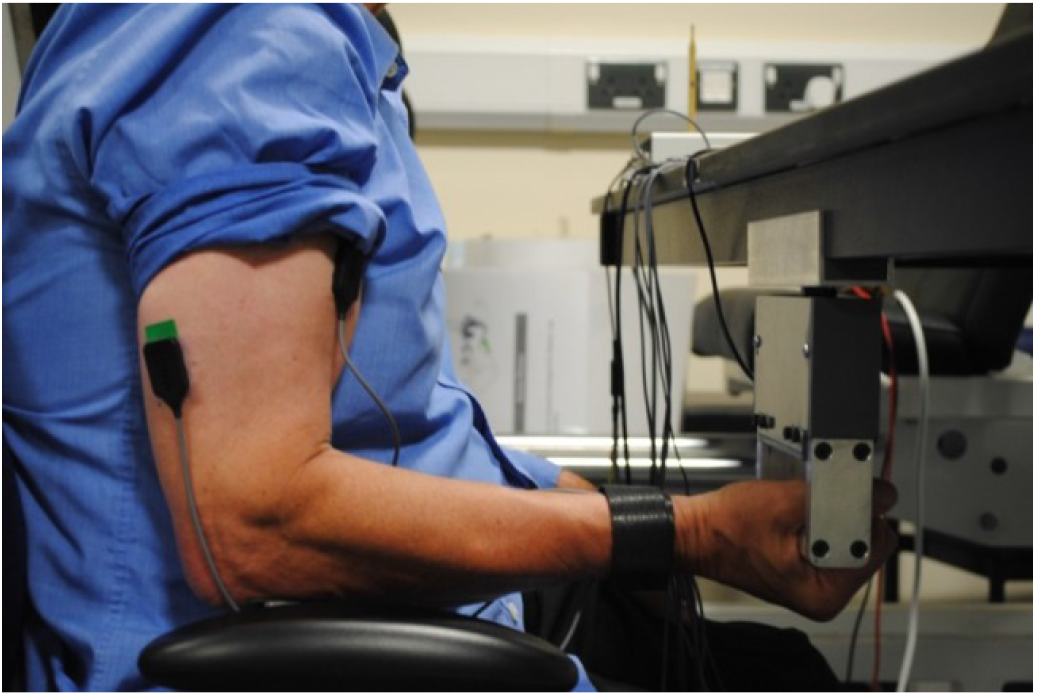
Photograph showing volunteer posture and sensor placement used for the tests on the ULV device

### Study design

Participants were asked to visit the laboratory 3 times (One visit to establish the MVC and familiarize with the equipment and 2 visits to receive the experimental treatment and assess its acute effects, see protocol in Figure 2).

**Figure 2:**
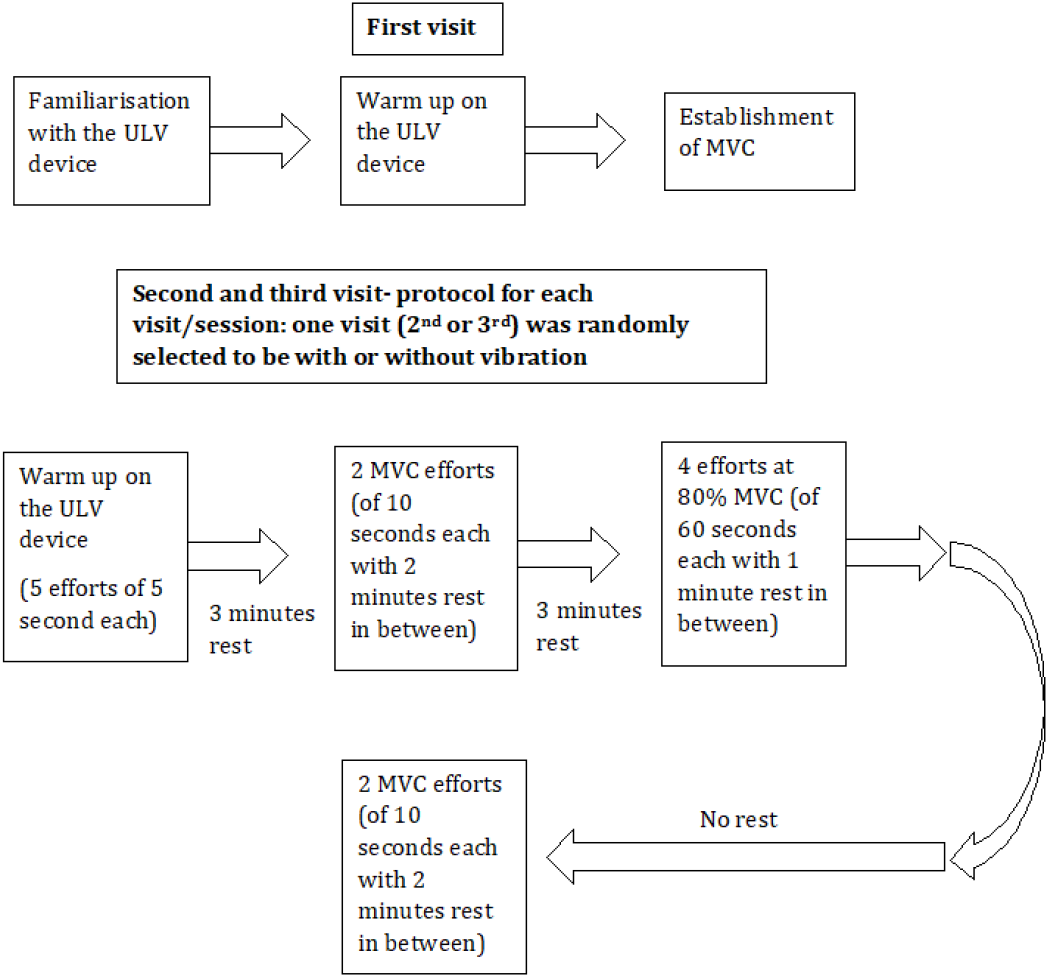
Schematic describing the arrangements/ research protocol of the experiment

In each laboratory visit the participants were fitted with the three-surface electromyography (sEMG) electrodes placed on the upper limb muscles: Biceps Brachii (BB), Triceps Brachii (TB), and Flexor Carpi Radialis (Forearm- FCR) muscle.

In the first visit each participant was introduced to the upper limb vibration (ULV) device (See Figure 1 and 2). Then the participant performed isometric arm flexion exercise of various intensities as a warm up by holding on to the hand grip of the ULV device with elbow flexed at 90°.

After the initial warm-up and familiarisation, the participants were asked to perform 5 Maximal Voluntary Contractions (MVC) performing an arm curl maximal effort for 10 seconds, each separated by a 5 minutes period of no-activity/resting. The average of the five efforts was used as the MVC value for that participant. In the second and third visits the participants went through the following vibration exercise treatment with a randomised cross-over design.

(A) Treatment 1: 4 sets of 1 minute muscle contraction (referred to as Control Fatiguing Contractions, i.e. CFC1, CFC2, CFC3 and CFC4) with 80% of MVC.

(B) Treatment 2: 4 sets of 1 minute muscle contraction (referred to as Vibration Fatiguing Contractions, i.e. VFC1, VFC2, VFC3 and VFC4) with 80% of MVC coupled with 30Hz vibration at 0.4mm amplitude.

The exercise treatments consisted of an isometric arm curl (flexion) against the ULV device hand-grip with the elbow flexed at 90° at the target force, for 60 seconds. 5 minutes’ rest was allowed between the two measurements. 8 measurements were carried out per session with a random selection, with a total of 16 measurements taken in the two sessions (second and third session).

Visual real time feedback from a load cell in the device and verbal feedback from the device operator were used to achieve the 80% of MVC force level. During each measurement, the sEMG activity of the designated muscles was recorded and stored for analysis along with the vibration characteristics and the force production details (load cell values). The vibration being delivered was continuously monitored with an accelerometer (ADXL-330, Analog Devices, MA, USA).

All the procedures were non-invasive. Medical grade, single use sEMG electrodes were used. Participants wore appropriate clothing to facilitate sensor placement on the upper limbs. Randomised cross over design was used to carry out these 16 exercise treatments/tests. As a cross over experiment, each participant was randomly allocated to one of the several possible sequences of the treatments.

### Instructions to the participants

Participants were asked and guided to maintain consistency in their hand grip positions and elbow angles. Both of these variables were measured continuously throughout the tests. Participants received both verbal and visual (real time graphical values on the PC) feedback to assist them in maintaining a constant force level.

### Fatigue and Safety

A minimum of 72 hours of recovery time was allowed between any two testing sessions to avoid any residue of delayed onset of muscle soreness (DOMS) and fatigue. Also, a general log of the participants’ daily physical activity excluding the vibrations tests was kept, i.e. participants who undertook any form of regular/irregular physical exercise e.g. strength or resistance training etc. Participants did not undergo the ULV exercise on the same day or immediately after finishing their regular exercise. At least a 1 day time gap was allowed between the regular physical exercise and ULV exercise, to avoid any effect of muscle fatigue (especially of upper limbs).

### EMG measurements and processing

sEMG was recorded from the BB, TB, and FCR during all exercise conditions according to recommendations reported in the literature [19]. Active bipolar electrodes (DelSys, Inc.; model: DE 2.1) were aligned with the muscle fiber direction and placed between the tendon and the muscle belly. To minimize the impedance and to ensure a proper contact, the skin was shaved as necessary, lightly abraded and cleaned with 70% isopropyl alcohol. The reference electrode was placed on an electrically inactive area of the lumbar spine (the anterior superior iliac spine). The sEMG electrodes and cables were secured to subject’s skin with medical tape. Active grounding and shielding of the cables was carried out to minimize electromagnetic inference [20]. The sEMG signals were sampled at 1000Hz, amplified with a gain of 1000 and analogue filtered for a 20–450Hz band pass with DelSys hardware (DelSys, Inc.; model: Bagnoli-4). Data acquisition was performed through a 16 bit data acquisition card (National Instruments Corp.; model: PCI-6220M) and EMGWorks (DelSys, Inc.) software.

Subsequent data processing and analysis was performed with custom written MATLAB code (The Mathworks, Inc.; version 8) routine. Any baseline offset of the sEMG data was removed by subtracting the mean.

The root mean square (RMS i.e. EMGrms) was used to estimate the neuromuscular activation. The RMS was calculated using the moving window technique. Initially the RMS was calculated for each window, and then the RMS for the entire data length was obtained by averaging the individual RMS values of each window.

The Mean Frequency (MEF) and Median Frequency (MDF) of the sEMG data was also obtained. These spectral estimators were also derived by moving window technique. Due to the similarity of trends shown by MEF and MDF, MDF results are excluded from any further discussion in this paper.

For both the amplitude and spectral estimation, the hamming window with a length of 1 second and no overlap was used. It has been shown that the choice of the window does not have a critical bearing on the spectral estimators like MEF and PSD [21]. Furthermore, for isometric, constant force and fatiguing contractions, the signal is regarded as stationary for epoch/window duration of 1 to 2 s. Previous studies suggest that epoch durations between 500 ms to 1 s provide better spectral estimation [21]–[23]. Also, it has been shown that window overlapping does not provide any significant benefits [21]. Based on these recommendations, the window length was kept to 1 s without any overlap.

The primary objective of this study was to quantify the fatiguing effect of ULV stimulation superimposed on isometric contraction on the targeted muscles. Due to these objectives, EMG data of the fatiguing contractions’ (Figures 3 and 2) was processed in a specific manner. These processing techniques are detailed below.

**Figure 3:**
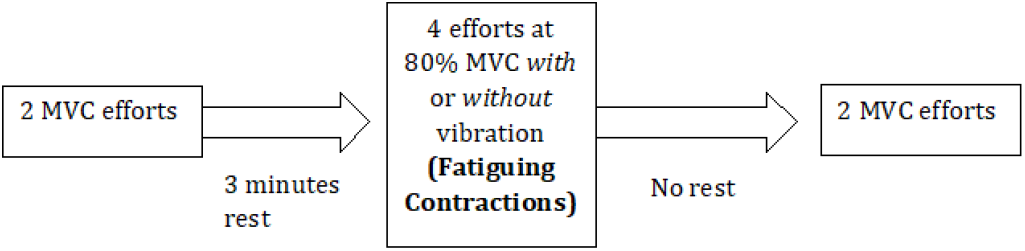
Schematic describing the MVC and Fatiguing Contractions

### EMG Data processing of the Fatiguing Contractions

During fatiguing contractions, different individuals performing similar relative effort (80% of their respective MVCs) are able to sustain the same level of force for a different period of time (for example, a highly trained young athlete would respond differently to the same fatigue task as compared to an elderly). Due to these differences, participants in this study sustained the same isometric fatiguing contractions for a different length of time. As described in the study design above, the target was to maintain the 80% of MVC force for 60 seconds under both control fatiguing contractions (CFCs) and vibration fatiguing contractions (VFCs). However, while certain individuals managed to maintain the target force for 60 seconds others did not. Especially as the treatment progressed from the first fatiguing effort to the second and so on (i.e. from CFC1 to CFC4 or from VFC1 to VFC4), the time duration for which an individual could maintain the target force dropped to 30 seconds in some cases (see Table 5) for the last fatiguing contraction (i.e. CFC4 or VFC4).

As the objective was to compare the relative fatigue levels attained by the individuals while performing the same relative efforts under vibration stimulation and control exercises, in order to have consistent data for comparisons, the EMG data selected for the analysis began once the target force level of 80% of MVC was achieved and was continued until the force level dropped down to 60% of MVC. This means the time for which the EMG data was selected was precisely connected to an individual’s ability to sustain the target force. As expected, this period varied with individual and with the progression to successive fatiguing contraction (Table 5).

This selected EMG data was then divided into five equal sections. Then an average for each section was calculated to give a single mean value for that section of the fatigue effort for that specific muscle under specific treatment condition. Both the time domain (EMGrms) and frequency domain (mean (MEF) and median frequency (MDF)) calculations were performed on the same data selection.

### Line Artifact Removal

Some authors have filtered the peaks in sEMG spectra coinciding with the vibration stimulation frequencies assuming them to be motion artifacts [24]. However it is still unclear whether the spectral peaks correlating with the stimulation frequencies are in fact motion artifacts [23] or stretch reflexes [13]. Recent evidence suggests that these peaks might indeed be stretch reflexes [25]. Considering the present ambiguity about the existence of motion artifacts and increasing evidence towards the presence of stretch reflex [13], [25], only the spectra exhibiting the largest power and hence potential to skew the results were removed. The largest spectra were found to be at 50Hz coinciding with mains frequency of some of the equipment. A Butterworth notch filter (10^th^ order, cut-off frequencies 49.5–50.5Hz) was employed to remove the components at this frequency.

### Statistical analysis

Normalization was performed by dividing the EMGrms of the entire section of the data value to be normalized by the maximum value obtained from the MVC effort of each participant. Alpha was set at 0.05. In each case, a significant difference was defined for a computed p-value ≤ 0.05. Paired student t-tests (one tail, different variance) were employed to compare the sEMG responses between the C and V conditions and to establish the significance level (P value) of the deviations from the means. The distribution of grouped data was assessed for normality using the Lilliefors test with a significance detection level of ≤ 0.05. Statistical analysis was carried out using the SigmaPlot statistical software package (Systat Software Inc.; Version SigmaPlot 12).

In the results below, statistically significant differences (P value ≤ 0.05) are noted by *.

## Results

### EMG Frequency Results- Pre and Post Fatigue Exercise-Mean frequency (MEF)

MVC efforts (MVC3 and MVC4) post both the vibration and control fatiguing treatments showed a reduction in MEF (and MDF) values in comparison with their respective pre-treatment effort values (MVC1 and MVC2), an expected effect of muscle fatigue manifested in myoelectric signal (Figures 4–6). Further, for MVC3 and MVC4, all the post vibration-treatment frequency values were higher than the respective post control-treatment values, this is unexpected. However only the Biceps’ MVC3 and MVC4 showed statistically significant higher values of MEF when control (C) and vibrations (V) values were compared. This is likely to be due to the Biceps being the prime mover in the elbow flexion exercise. Higher MEF values of post vibration treatment compared to the control are slightly puzzling as one would expect higher fatigue manifestation (post vibration) hence lower MEF and MDF values compared to the control. These unexpected MEF values could be the effect of removing 50Hz frequencies from the EMG signal leading to a shift in the spectral compression or this change could be due to potential effect of recruitment of large MUs or MU synchronisation.

**Figure 4:**
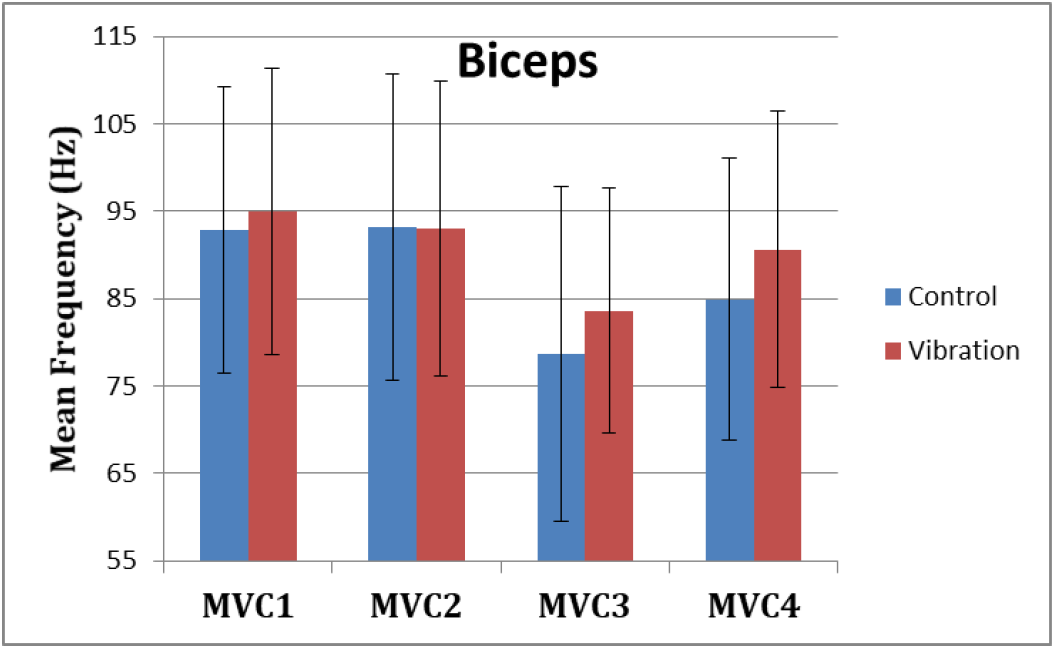
Mean frequency (MEF) values for the Biceps during each MVC effort pre (MVC1 and MVC2) and post (MV3 and MVC4) fatigue exercise under control (no vibration) and vibration (30Hz-0.4mm) conditions.

**Figure 5:**
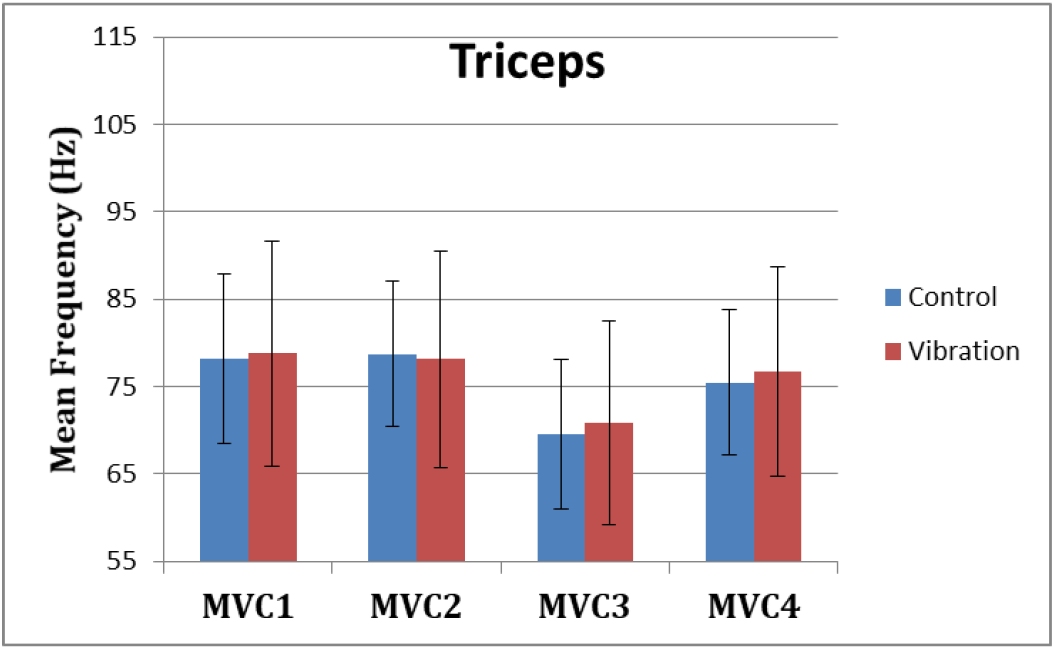
MEF values for the Triceps during each MVC effort pre (MVC1 and MVC2) and post (MV3 and MVC4) fatigue exercise under control (no vibration) and vibration (30Hz-0.4mm) conditions.

**Figure 6:**
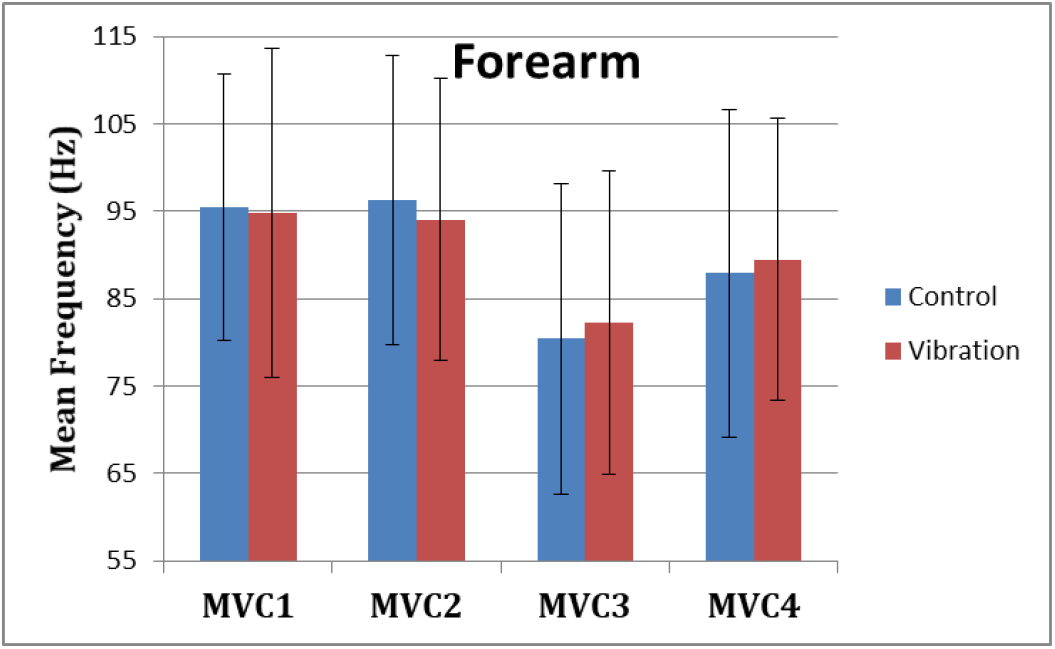
MEF values for the Forearm during each MVC effort pre (MVC1 and MVC2) and post (MV3 and MVC4) fatigue exercise under control (no vibration) and vibration (30Hz-0.4mm) conditions.

Comparison between pre and post fatiguing contraction MVCs for the vibration treatment (vibration MVC2 and MVC3) resulted in statistically non-significant difference whereas a similar comparison of the control treatment efforts indicates statistically significant differences in all the muscle groups (Table 1 and 2). These differences between the post-control and vibration treatment could be due to alteration in the fatigue mechanisms through the CNS, such as potentiation of stretch reflex, to cope with the higher neuromuscular demand induced by vibration stimulation. Also, the vibration MEF values for MVC4 for all the muscles groups are higher compared to corresponding control MVC4 values, suggesting continued effect of vibration treatment post recovery period of 3 minutes. Although none of the vibration (MVC4) values were (statistically) significantly higher than their respective controls, this result indicates that the vibration treatment superimposed with isometric contraction alters neuromuscular strategy post fatiguing conditions.

**Table 1:** Comparison between MEF values of each control and vibration effort showing statistical significance (^*^).

|  | MVC1 | MVC2 | MVC3 | MVC4 |
| --- | --- | --- | --- | --- |
| Biceps | P value = 0.227 | P value = 0.231 | P value = 0.034* | P value = 0.029* |
| Triceps | P value = 0.429 | P value = 0.183 | P value = 0.298 | P value = 0.321 |
| Forearm | P value = 0.408 | P value = 0.124 | P value = 0.236 | P value = 0.333 |

**Table 2:** Comparison between MEF values of effort performed just before (MVC2) and after (MVC3) the fatiguing exercise, showing statistical significance (^*^).

|  | Comparison between Control MVC2 and MVC3 | Comparison between Vibration MVC2 and MVC3 |
| --- | --- | --- |
| Biceps | P value = 0.001* | P value = 0.414 |
| Triceps | P value = 0.001* | P value = 0.413 |
| Forearm | P value = 0.002* | P value = 0.334 |

### EMG Amplitude Results- Pre and Post Fatigue Exercise-EMGrms

The MVC efforts post both the vibration and control fatiguing treatments (MVC3 and MVC4) resulted in an increase in mean EMGrms amplitudes in comparison with their respective pre-treatment effort values (MVC1 and MVC2). Further, within the increased post treatment effort EMGrms values i.e. MVC3 and MVC4, Biceps (MVC3, MVC4) and Triceps (MVC3) show statistically significant decrease in EMGrms post vibration treatment, compared to the control. This decrease could be due to the mechanical manifestation of higher level of fatigue induced by the vibration treatment compared to the control. Comparison between the pre and post fatiguing contraction MVCs (MVC2 and MVC3) of the controls result in statistically significant increases in EMGrms of all the muscles. Whereas, comparison between the pre and post fatiguing contractions of vibration MVCs (MVC2 and MVC3) result in non-significant differences in all the muscle groups. This further suggests inability of the muscles to produce the similar levels of activation as the controls, post-vibration treatments. This implies a higher level fatiguing effect on the muscles due to enhanced neuromuscular demand put on them by vibration superimposed on isometric contractions.

**Table 3:** Comparison between normalised EMGrms values of each control and vibration effort showing statistical significance.

|  | MVC1 | MVC2 | MVC3 | MVC4 |
| --- | --- | --- | --- | --- |
| Biceps | P value = 0.035* | P value = 0.060 | P value = 0.191* | P value = 0.371* |
| Triceps | P value = 0.008* | P value = 0.036* | P value = 0.045* | P value = 0.171 |
| Forearm | P value = 0.166 | P value = 0.139 | P value = 0.484 | P value = 0.141 |

**Table 4:** Comparison between normalised EMGrms values of effort performed just before (MVC2) and after (MVC3) the fatiguing exercise, showing statistical significance.

|  | Comparison between Control MVC2 and MVC3 | Comparison between Vibration MVC2 and MVC3 |
| --- | --- | --- |
| Biceps | P value = 0.024* | P value = 0.025 |
| Triceps | P value = 0.027* | P value = 0.026 |
| Forearm | P value = 0.256* | P value = 0.059 |

### EMG Frequency Results- During Fatigue Exercise- Mean Frequency (MEF)

Within each fatiguing effort i.e. CFC or VFC, as the exercise/treatment progressed in time (i.e. consecutive sections of the fatigue tasks), the MEF’s trend under vibration conditions separated from that of the control conditions. The MEF under vibration condition efforts decreased with a steeper slope indicating a higher rate of fatigue manifestation compared to the corresponding control effort (Figures 10–17). The separation between vibration and control mean frequencies is more evident for the Triceps and Forearm muscles than the Biceps (Figures 10–17).

As the consecutive fatiguing efforts progressed i.e. from CFC1, CFC2, CFC3 to CFC4 and for VFC1 to VFC4), the separation between vibration and control MEF increased. Thus MEF values under vibration condition decrease further in value in comparison with their control counterpart at the last effort (CFC4 vs. VFC4) as compared to their separation in the initial efforts (e.g. CFC1 vs. VFC1). This indicates, as the exercise efforts progressed, vibration stimulation combined with isometric contraction continued to induce higher neuromuscular demand leading to increased fatigue levels in the engaged muscles compared to the control exercises. Also, participants were able to sustain the fatiguing contractions for less required times under the vibration compared to the control (Table 5). This further indicates higher fatigue manifestation under the vibration conditions compared to the control. However, only Triceps and Forearm showed statistically significant lower MEF values under vibration conditions as compared to their respective control conditions. This trend is puzzling as the Biceps is the main prime mover muscle in this exercise and one would expect more fatigue hence higher separation between the vibration and control conditions’ MEF values in this muscle. Nevertheless, all the three muscles showed trends of decrease in frequencies described above. This higher rate of fatigue under vibration superimposed on contraction could be due to the changes in MU recruitment patterns. It is known that recruitment of the smaller MU leads to reduction in MEFs, as these MUs have smaller conduction velocity (leading to lower MEF) [26]. As the exercise progressed to the next consecutive effort i.e. from CFC1 to CFC2 and so on, gradual decrease in the overall MEF can be observed irrespective of the (vibration/control) treatment condition and the muscle group (Figures 10–17). For example average MEFs in the second contraction effort (CFC2) are lower than the first (CFC1), possibly an effect and indication of muscle fatigue.

**Table 5:** Average times (in seconds) completed by participants during fatiguing contractions and their standard deviations.

| Control | CFC1 | CFC2 | CFC3 | CFC4 |
| --- | --- | --- | --- | --- |
| Times (s) | 53 | 51.30 | 44.53 | 40.53 |
| SD | 8.87 | 11.96 | 14.17 | 17.03 |
| Vibration | VFC1 | VFC2 | VFC3 | VFC4 |
| Times (s) | 49.92 | 49.23 | 43.50 | 40.53 |
| SD | 13.18 | 12.79 | 14.83 | 16.95 |

### EMG Amplitude Results- During Fatigue Exercise-EMGrms

Within each fatiguing effort i.e. CFC or VFC, as the exercise/treatment progressed in time, the EMGrms amplitude trend under vibration conditions separated more from the control conditions. EMG amplitude under vibration condition efforts increased with a steeper positive slope indicating higher neuromuscular activity compared to the corresponding control effort (Refer supplementary information, Figures 18–29). As the consecutive fatiguing efforts progressed the separation between the vibration and control EMG amplitudes mostly remained at the same level (Figures 18–29). Thus the EMGrms amplitude values under vibration condition were higher (with the exception of Triceps CFC4 vs. VFC4) compared with their control counterpart despite the progression of fatigue tasks from CFC1/VFC1 to CFC4/VFC4. This suggests, that as the exercise efforts progressed, the vibration stimulation combined with isometric contraction induced higher neuromuscular activity leading to increased EMG amplitude levels in the engaged muscles compared to the control exercises. However, only the Forearm showed statistically significantly higher EMGrms amplitude values under vibration conditions compared to the respective control conditions (Figures 26–29). Nevertheless, all three muscles showed trends of EMG amplitude separation described above with the Forearm showing the largest difference between the vibration and the control treatment EMGrms values followed by Biceps and Triceps.

## Discussion

### Overall Fatiguing Effects

The effectiveness and the ability of vibration exercise(s) to induce enhanced neuromuscular activity have been investigated primarily through EMG amplitude responses. Only a small number of studies have investigated the fatiguing effects of vibration stimulation through the analysis of EMG frequency variables. Mischi et al. and Xu et al. have recently investigated the fatiguing effects of vibration through sEMG frequency analysis [15], [27]. Our study design and the vibration stimulation device differs from these earlier reported studies [15], [27].

The results of this ULV study confirm that in comparison with the near-maximal (80% of MVC) isometric contraction alone, near-maximal (80% of MVC) isometric contraction superimposed on vibration stimulation (of 30Hz-0.4mm) elicits an increased neuromuscular response in the upper limbs. This leads to a higher fatigue in the engaged muscles during the vibration treatment, as judged through decreased capacity to maintain specific force levels. Equally importantly, the results also indicate that muscles display a higher degree of myoelectric manifestation of fatigue during vibration treatment efforts compared to the control. The results also imply potential alterations in neuromuscular mechanisms to cope with the higher neuromuscular load/demand induced by the vibration superimposed on isometric contraction.

Although all three muscles investigated display higher fatigue and enhanced neuromuscular activity, the effect of vibration stimulation is different on different muscles. It was anticipated that due to their role in flexion, the agonist muscles (Biceps and Forearm) would exhibit higher fatigue and neuromuscular activity than the antagonist (Triceps). The results indeed confirm that the Biceps and Forearm display higher neuromuscular activity (EMGrms) during vibration treatment/fatigue efforts over the Triceps muscles. However, interestingly the Triceps also displayed more significant (P < 0.05) reduction in MEF, hence higher fatigue levels compared to Biceps during the vibration treatment/fatigue efforts. Co-activation of the agonist and antagonist has been reported to maintain the joint angle/stability during the vibration stimulation in the upper limbs [28]. It seems that although the Triceps is not the prime mover in the force producing flexion task, its role in maintaining the elbow joint stability during the stimulation has effect on its neuromuscular activity leading to lower MEF, hence higher fatigue.

The difference in the neuromuscular response of the muscles could be attributed partially to their proximity to the vibration source. A higher vibration transmission can lead to comparably higher neuromuscular activity over the corresponding control treatment. The Forearm was the most proximal muscle group to the vibration actuator being investigated here and show the most significant (P < 0.05) higher neuromuscular response (EMGrms) and the most significant (P < 0.05) fatigue manifestation (MEF and MDF) compared to the control condition during the vibration treatment/fatigue tasks.

The difference in the neuromuscular response of the muscles to the vibration stimulation also depends on the muscle pre-stretch i.e. muscle length/contraction prior to the stimulation along with the vibration characteristics [29], [30]. This might partially explain why despite showing relatively higher neuromuscular activity and fatigue manifestation during the vibration treatment/efforts (Figures 15–17), the Forearm did not display a similar level of fatigue manifestation post treatment as the Biceps (Figures 4 and 6, Table 1). The Biceps displayed the most significantly (P < 0.05) different EMGrms values post vibration treatment (MVC3 and MVC4) compared to the control (Figures 7–9, Table 3). The higher level of fatigue manifestation in Biceps post vibration treatment than the Forearm could be also related to the higher level of pre-contraction the Biceps had to produce in order to facilitate the elbow flexion task. The results strongly indicate that vibration stimulation due to its higher neuromuscular demand potentially alters the underlying fatigue mechanisms. These potential fatigue mechanisms are discussed in the section below.

**Figure 7:**
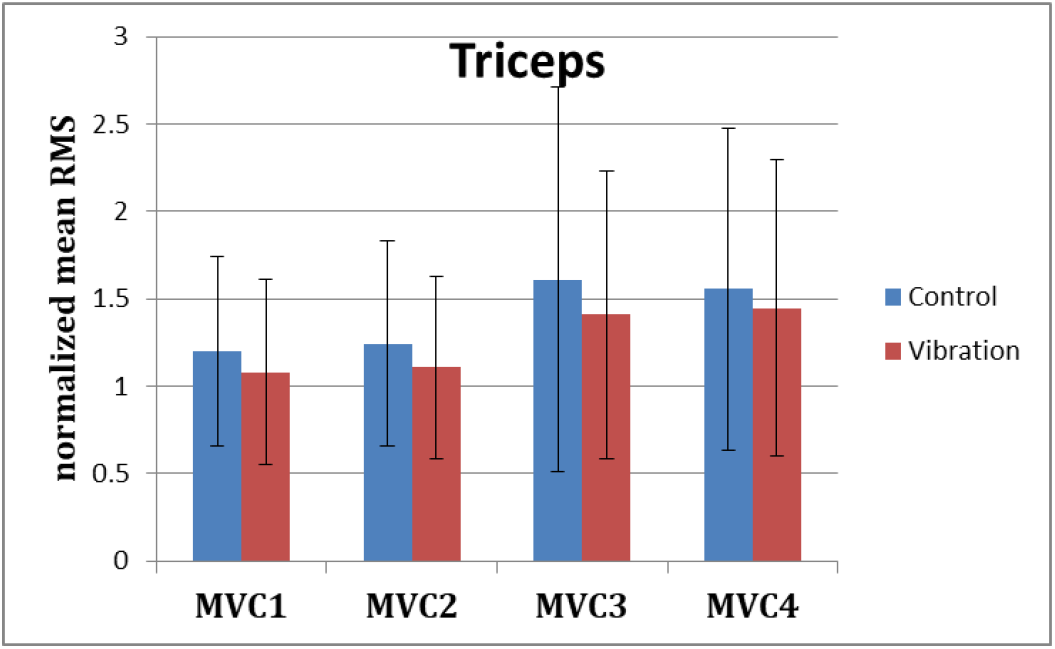
Normalised EMGrms values for the Biceps during each MVC effort pre (MVC1 and MVC2) and post (MV3 and MVC4) fatigue exercise under control (no vibration) and vibration (30Hz-0.4mm) conditions.

**Figure 8:**
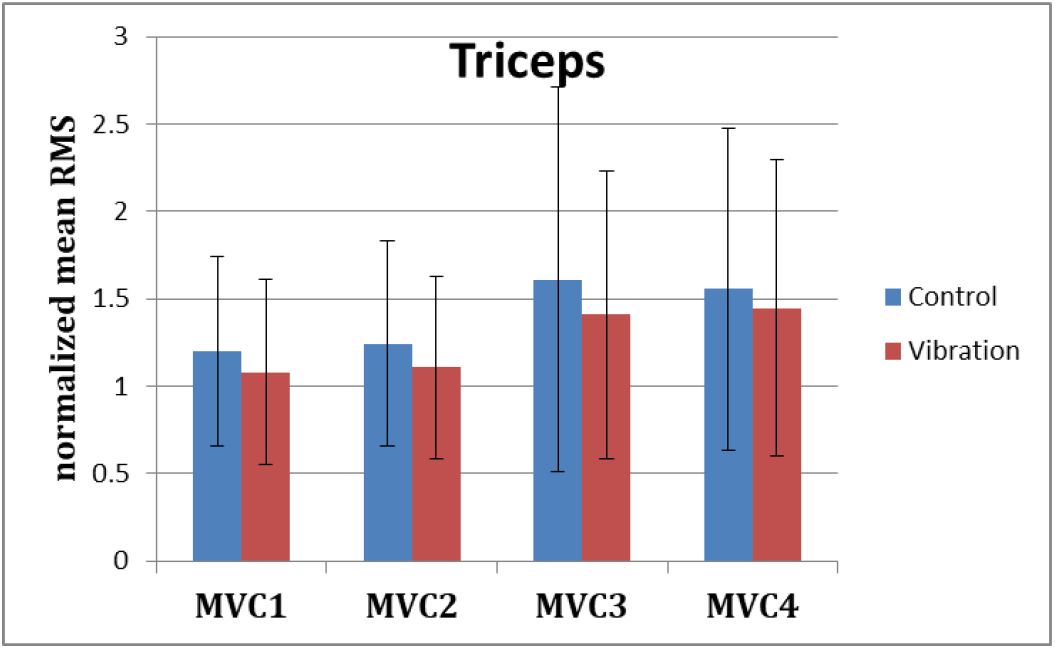
Normalised EMGrms values for the Triceps during each MVC effort pre (MVC1 and MVC2) and post (MV3 and MVC4) fatigue exercise under control (no vibration) and vibration (30Hz-0.4mm) conditions.

**Figure 9:**
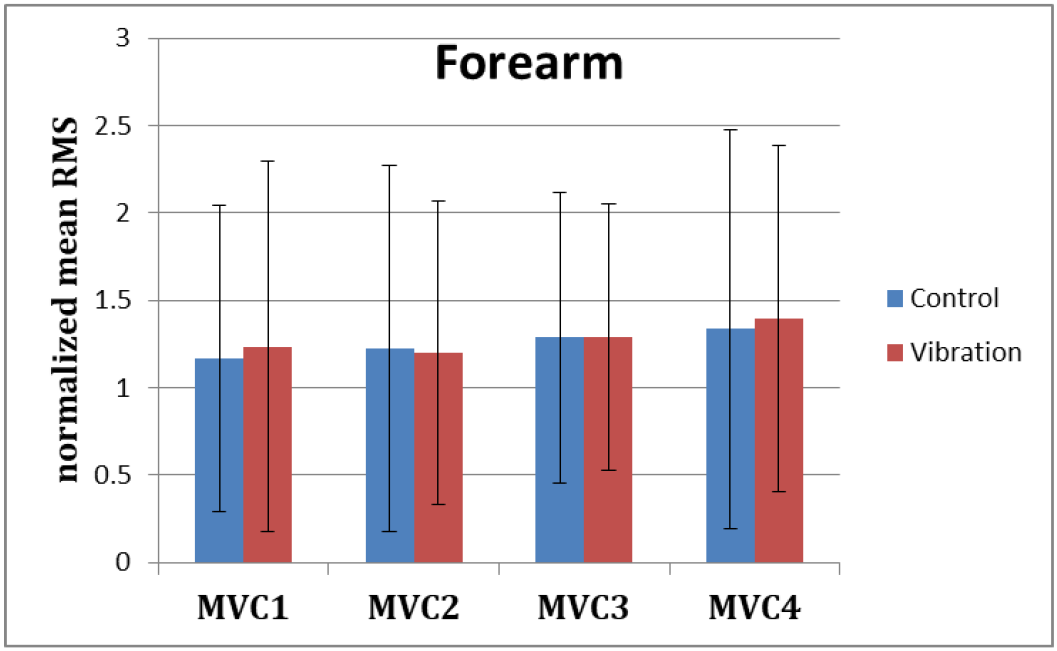
Normalised EMGrms values for the Forearm during each MVC effort pre (MVC1 and MVC2) and post (MV3 and MVC4) fatigue exercise under control (no vibration) and vibration (30Hz-0.4mm) conditions.

**Figure 10:**
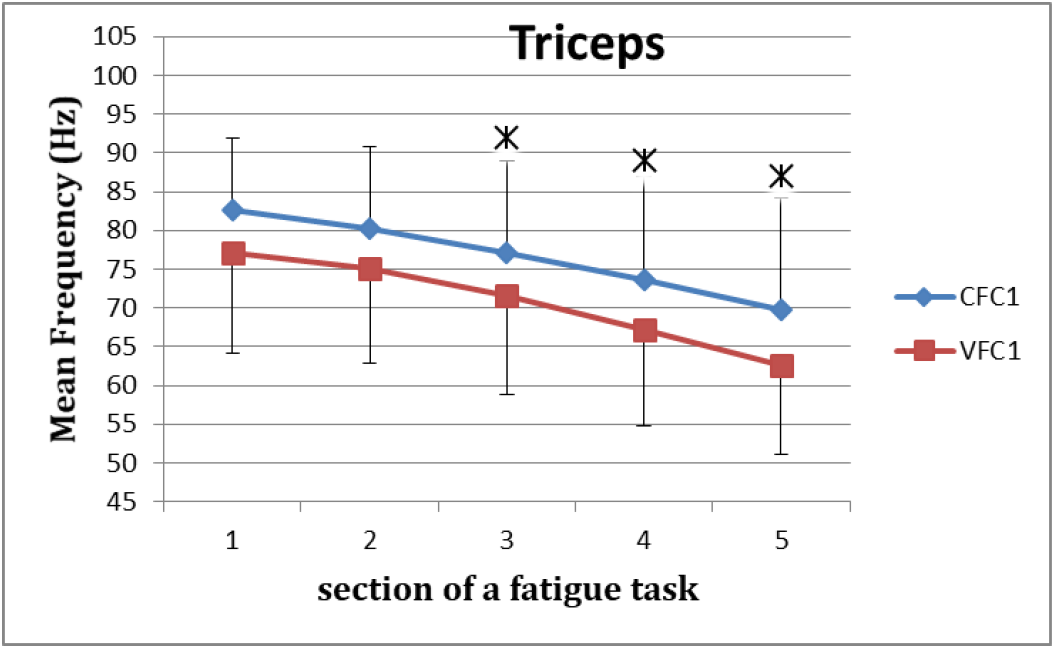
MEF values for the Triceps for the five consecutive sections of the fatigue effort during the progression of each of the successive four fatiguing exercise efforts performed, under control and vibration condition, CFC1 vs VCF1

**Figure 11:**
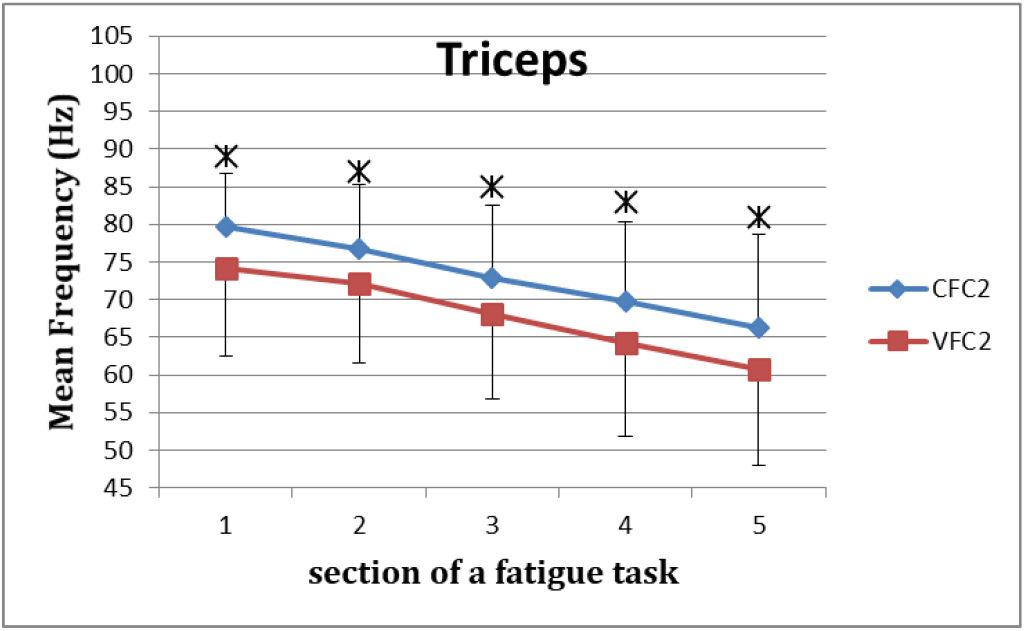
MEF values for the Triceps for the five consecutive sections of the fatigue effort during the progression of each of the successive four fatiguing exercise efforts performed, under control and vibration condition, CFC2 vs VCF2

**Figure 12:**
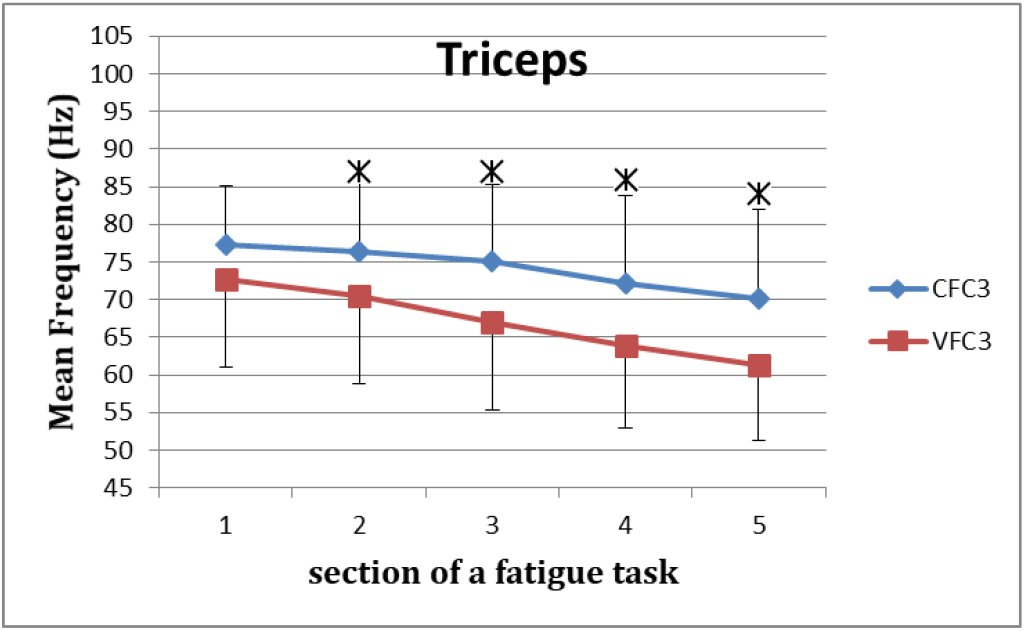
MEF values for the Triceps for the five consecutive sections of the fatigue effort during the progression of each of the successive four fatiguing exercise efforts performed, under control and vibration condition, CFC3 vs VCF3

**Figure 13:**
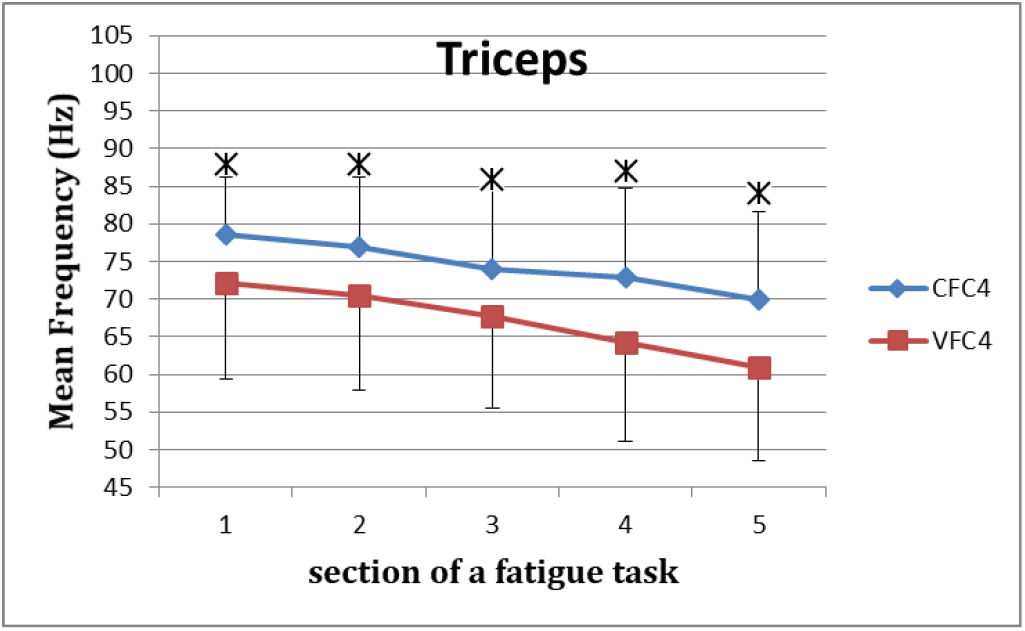
MEF values for the Triceps for the five consecutive sections of the fatigue effort during the progression of each of the successive four fatiguing exercise efforts performed, under control and vibration condition, CFC4 vs VCF4

**Figure 14:**
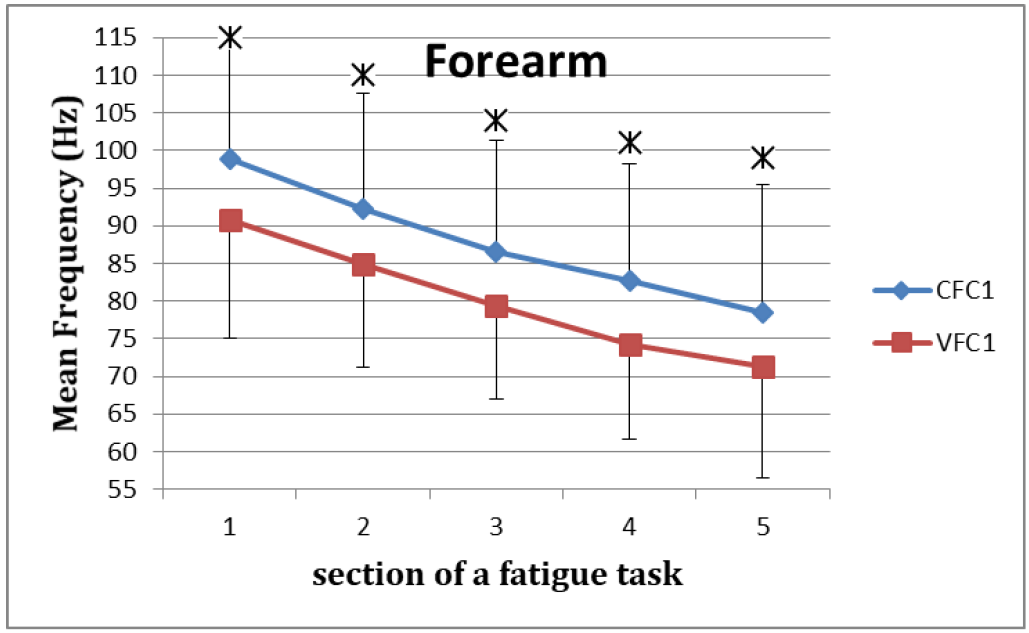
MEF values for the Forearm for the five consecutive sections of the fatigue effort during the progression of each of the successive four fatiguing exercise efforts performed, under control and vibration condition, CFC1 vs VCF1

**Figure 15:**
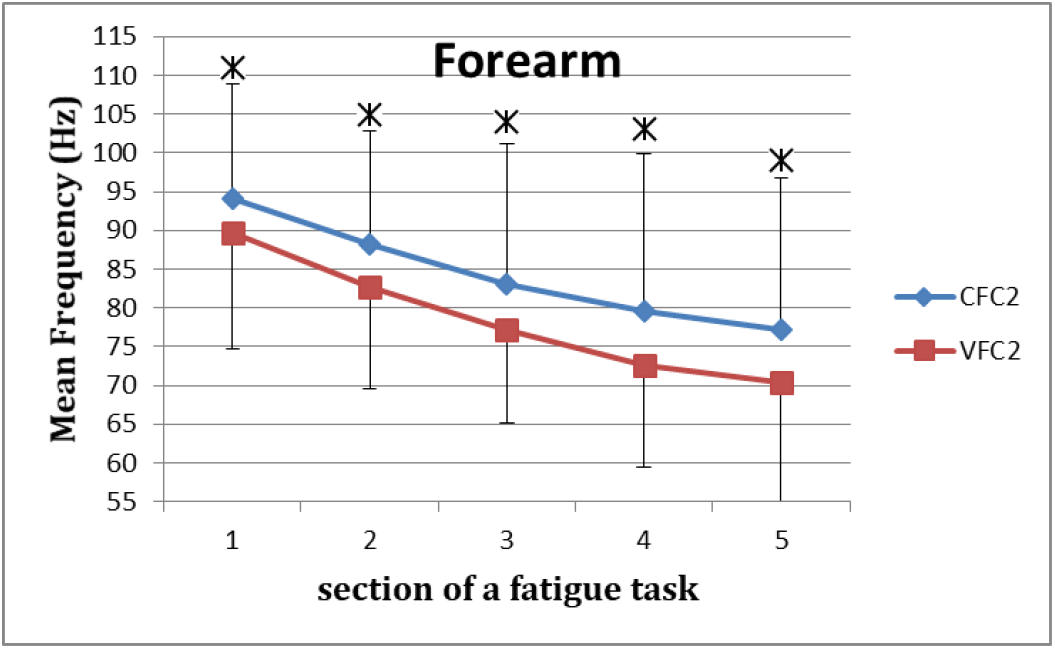
MEF values for the Forearm for the five consecutive sections of the fatigue effort during the progression of each of the successive four fatiguing exercise efforts performed, under control and vibration condition, CFC2 vs VCF2

**Figure 16:**
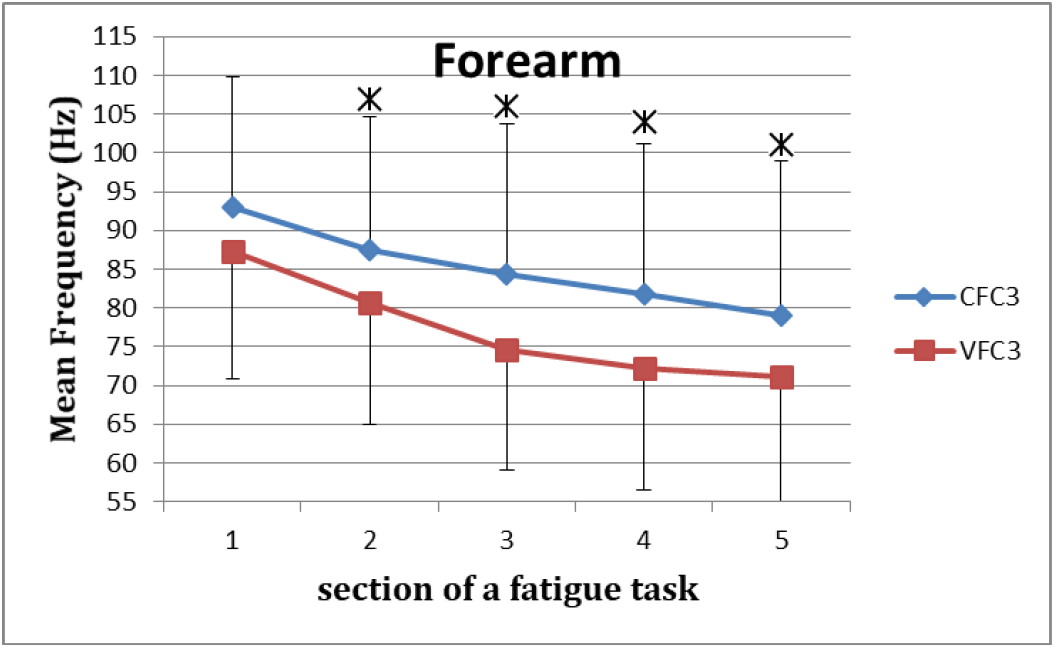
MEF values for the Forearm for the five consecutive sections of the fatigue effort during the progression of each of the successive four fatiguing exercise efforts performed, under control and vibration condition, CFC3 vs VCF3

**Figure 17:**
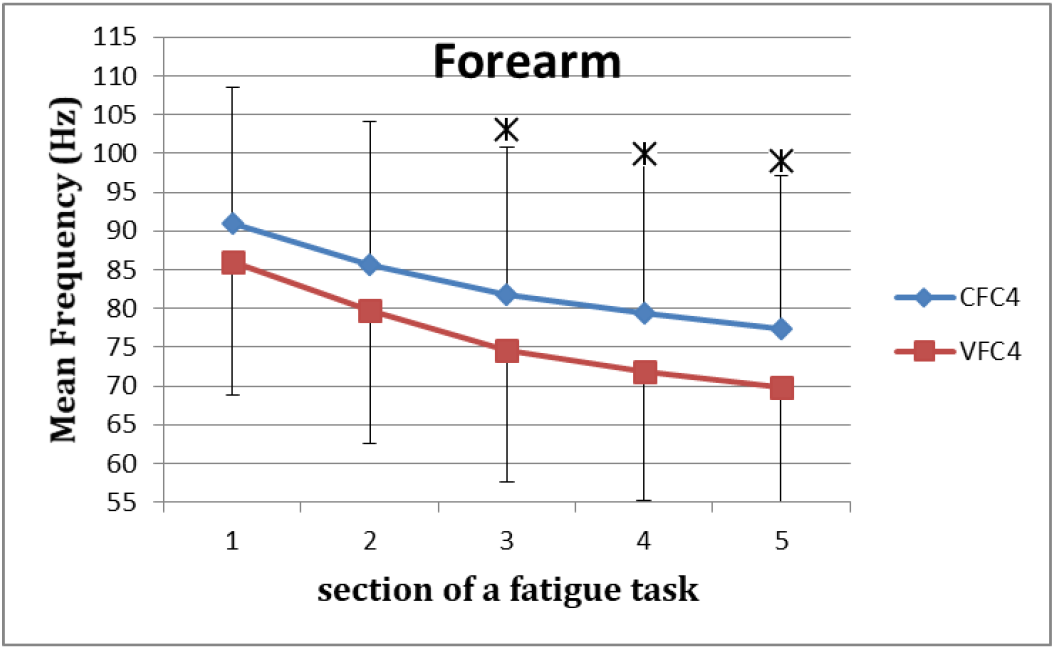
MEF values for the Forearm for the five consecutive sections of the fatigue effort during the progression of each of the successive four fatiguing exercise efforts performed, under control and vibration condition, CFC4 vs VCF4

### Potential Mechanisms leading to Increased Neuromuscular Activity and Fatigue

It has been reported that myoelectric manifestations of fatigue are induced by the two main physiological factors, peripheral (muscle) fatigue and central fatigue [31]. The peripheral/muscle fatigue relates to the reduction of conduction velocity of MU action potential and the central fatigue is characterised by the synchronisation of MUs by the CNS to increase the mechanical output when the MU pool is recruited [22], [31]. Based on the above relation it has been suggested that the conduction velocity, MEF (MDF) values of the sEMG are strongly indicative of the peripheral fatigue [17]. With regards to the exercises in this study, during isometric constant force fatiguing contractions, conduction velocity seems to be particularly correlated with MEF of the sEMG [32]. Thus one can argue, MEF results in this study can reliably reflect CV and hence the peripheral fatigue.

Further, it is well established that during sustained contractions the power spectral density of the sEMG signal compresses and shifts towards the lower frequencies leading to a reduction in MEF/MDF [33], [34]. MEF values in this study have shown clear and consistent reduction as the fatiguing efforts progress and the MEF under vibration have shown higher reduction in MEF, which is indicative of reducing CV in active motor units. Further, decrease in CV is related to increase in fatigue [33], [34]. Based on the higher reduction MEF values compared to the control, the results of this study indicate strongly that vibration superimposed on isometric contraction induces higher fatigue compared to the control. This was true of all the muscles tested in this study, that is agonist (Biceps and Forearm) and antagonist (Triceps). Further, all the above muscles have also shown higher EMGrms values, corresponding to their lower MEF values, in almost all the vibration conditions compared to the control. Increase in EMGrms have been associated with increase in fatigue under sustained contractions [35]. This further affirms the higher fatiguing effects of vibration exercise superimposed with isometric contractions.

It is important to note that the vibration superimposed with isometric contractions continued to induce higher rate of fatigue as the exercise progressed to the subsequent efforts, thus accentuating the overall fatiguing effect in the engaged muscles compared to their respective control conditions.

The enhanced neuromuscular activity observed during vibration stimulation has been also ascribed to the tonic vibration reflex (TVR). Further, although not in indirect vibration settings (i.e. WBV/ULV), however in direct vibration settings of upper limbs, increase in muscle fatigue has been attributed to the TVR [36]. It has been reported that MU synchronisation has direct effect on the TVR and hence the muscle fatigue [36], [37]. Thus the MU synchronisation and hence the central fatigue mechanisms could play a role in the responses to vibration stimulation superimposed with isometric contractions, although the exact mechanisms of MU synchronisation and central fatigue under indirect vibration stimulations are still unclear. Given that vibration stimulation can change the central motor commands excitability, CNS’s role in influencing responses to vibration superimposed with isometric contractions cannot be excluded.

Further it is important to note that, during MVC immediately post vibration treatment (i.e. MVC3) all the muscles displayed significantly higher MEF (and MDF) values respective to their controls and higher values during MVC after the recovery (i.e. MVC4). This is puzzling and one would expect the opposite, i.e. lower MEF (and MDF) values post vibration treatment. This is despite their corresponding EMGrms values being lower, hence indicating higher mechanical fatigue post vibration treatment compared to the control. The increased MEF (and MDF) values could be due to potentiation of the stretch reflex observed following the indirect vibration exercise, which leads to the recruitment of high threshold MUs leading to the increase in MEF (and MDF) [38].

To summarise, the above results confirm that vibration superimposed on isometric contraction not only leads to higher fatigue levels in the engaged muscles but also alters the neuromuscular function immediately post treatment.

### Limitations of the Study

The vibration characteristics of this study were based on previous evidence which indicated that 30Hz frequency stimulation combined with near maximal (> 70% MVC) isometric contraction could elicit the largest neuromuscular response compared to control condition in upper limbs [15]. Based on this evidence, the experiments in this study were limited only to the specific variables of excitation, i.e. 30Hz-0.4mm, 80% of MVC force level, Biceps, Forearm and Triceps muscles in elbow (isometric) flexion task. Hence the fatiguing effects of the stimulation and conclusions drawn from this study should not be extrapolated to other vibration characteristics, contraction types and levels and muscle groups.

In this study, no filtering was performed to remove the so called ‘motion artifacts’ from the sEMG data obtained. This might be considered a limitation. However it is important to note that a recent ULV study found no differences in the MEF, MDF and RMS values after removing the so called motion artifacts [27]. Further, another study by the same group have reported that the so called ‘motion artifacts’ are in fact ‘stretch reflex’ responses and hence filtering them might actually remove important information about stretch reflex responses and central fatigue mechanisms [25].

## Conclusions

(A) Near maximal (80% of MVC) isometric fatiguing contractions superimposed on vibration stimulation lead to a higher rate of fatigue development compared to the isometric contraction alone in the upper limb muscles.
(B) The higher rates of fatigue observed under vibration in all the muscles studied are associated with correspondingly higher rates of neuromuscular activations.
(C) Vibration superimposed on isometric contraction, continues to induce higher fatigue levels in the subsequent fatiguing efforts thus accentuating the overall fatiguing effect on the engaged muscles.
(D) Post vibration treatment MVCs show, higher manifestation of mechanical fatigue compared to the control. However, these post vibration MVCs also display higher levels of MEF (and MDF) implying alteration in neuromuscular function post treatment.
(E) Vibration superimposed on isometric contraction not only seems to alter the neuromuscular function during fatiguing efforts by inducing higher neuromuscular load but also post vibration treatment, potentially through the augmentation of stretch reflex and/or higher central motor command excitability.
(F) Both peripheral and central fatigue mechanisms are likely to play role in the alteration of neuromuscular function observed, however the role of peripheral fatigue mechanisms may be more prominent.
(G) The observed increase in fatigue and neuromuscular activity under vibration conditions are evident not only in agonist but also antagonist muscles, although the effect of vibration seem to depend on the muscle role (agonist/antagonist), muscle pre-stretch length, contraction level, vicinity to the vibration source and the vibration parameters.

## Acknowledgment

Authors would like to thank Scottish Funding Council (SFC) for the North East of Scotland Technology (NESTech) Seed Fund to support the study.

